## Supplementary Figures for "Epigenetic memory is governed by an effector recruitment specificity toggle in Heterochromatin Protein 1"

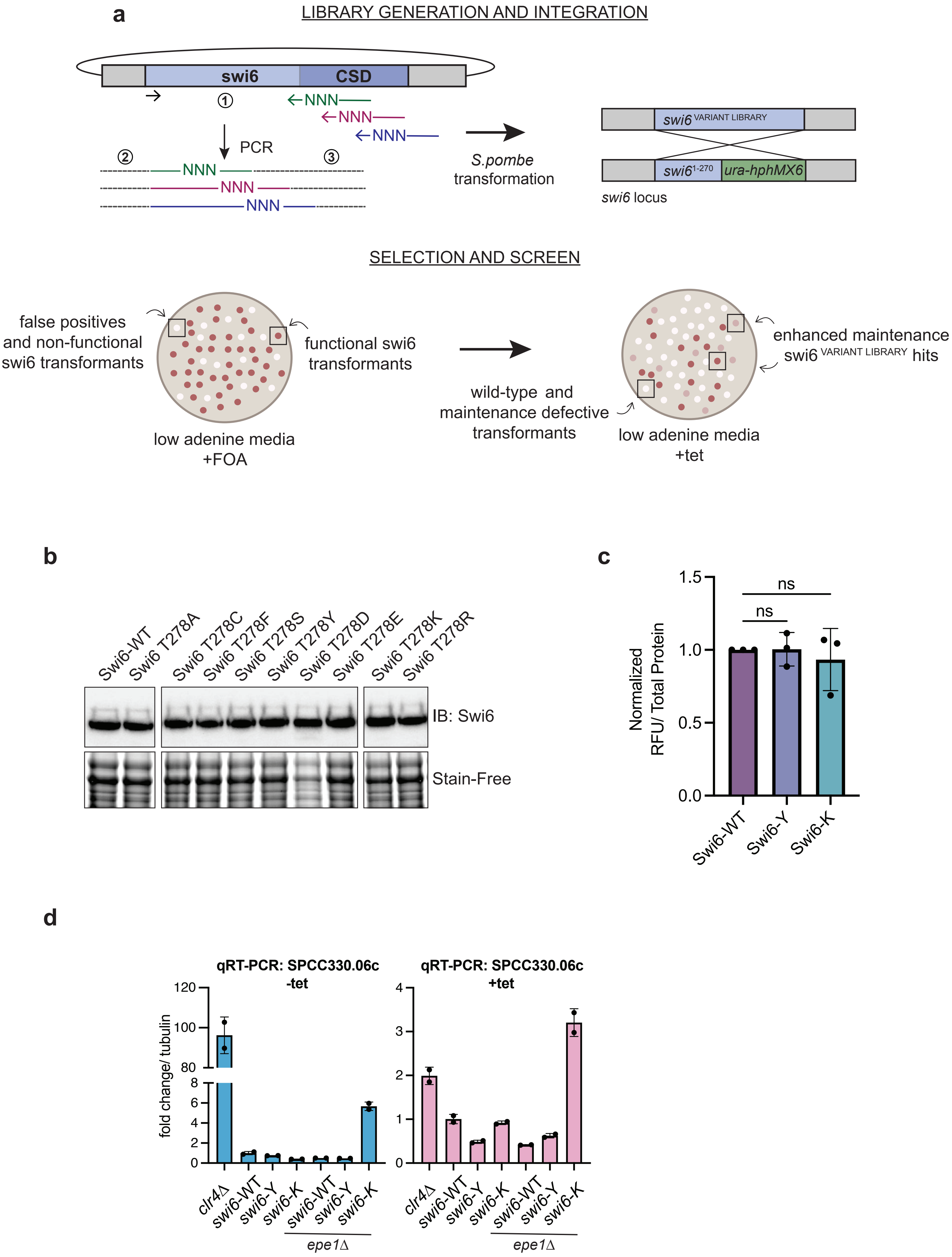
a
LIBRARY GENERATION AND INTEGRATION
swi6
CSD
NNN
PCR
S.pombe transformation
swi6 VARIANT LIBRARY
swi6 1-270
ura-hphMX6
swi6 locus
SELECTION AND SCREEN
false positives and non-functional swi6 transformants
functional swi6 transformants
low adenine media +FOA
wild-type and maintenance defective transformants
low adenine media +tet
enhanced maintenance swi6 VARIANT LIBRARY hits
b
Swi6-WT
Swi6 T278A
Swi6 T278C
Swi6 T278F
Swi6 T278S
Swi6 T278Y
Swi6 T278D
Swi6 T278E
Swi6 T278K
Swi6 T278R
IB: Swi6
Stain-Free
c
Normalized RFU/ Total Protein
ns
ns
Swi6-WT
Swi6-Y
Swi6-K
d
qRT-PCR: SPCC330.06c -tet
qRT-PCR: SPCC330.06c +tet
fold change/ tubulin
clr4Δ
swi6-WT
swi6-Y
swi6-K
epe1Δ

**Supplementary Figure 1. A targeted mutagenesis approach to identify variants within the chromoshadow domain that selectively influence heterochromatin maintenance.**
(**a**) Schematic outlining the technical design of our 3-step PCR-based site-directed targeted mutagenesis library approach. 1) A plasmid containing *swi6* with homology is used a template for PCR using a universal forward primer and reverse tiling primers containing a degenerate "NNN" codon targeting each amino acid position. 2) PCR product acts as a unidirectional primer to create longer swi6 products. 3) Amplify homologous flanking sequences for integration at the swi6 locus, where it can replace a nonfunctional *swi6-ura4-hphMX6* transgene. Clones were selected for *ade6+* silencing (cells grow red) on low adenine media, survival on FOA, and loss of hygromycin resistance. (**b**) Western Blot measuring Swi6 protein levels of Swi6-WT and Swi6 Thr 278 variants. Swi6 is detected using a Swi6 primary antibody and total protein is detected using stain-free imaging. (**c**) Protein expression of Swi6-WT, Swi6-Y, and Swi6-K quantified from a western blot using a fluorescent-labeled secondary antibody. Total protein was measured using Stain-Free imaging and fluorescence signal was measured using Alexa 647 detection. Images were analyzed using ImageJ and an unpaired two-tailed t-test was performed (P-value cutoff < 0.05). Error bars indicate SD (N=3). (D) qRT-PCR measuring RNA levels at *SPCC330.06c* in indicated genotypes before (-tet) and after (+tet) tetracycline treatment. Error bars indicate SD (N=2). (**d**) qRT-PCR measuring RNA levels at *SPCC330.06c* in indicated genotypes before (-tet) and after (+tet) tetracycline treatment. Error bars indicate SD (N=2).

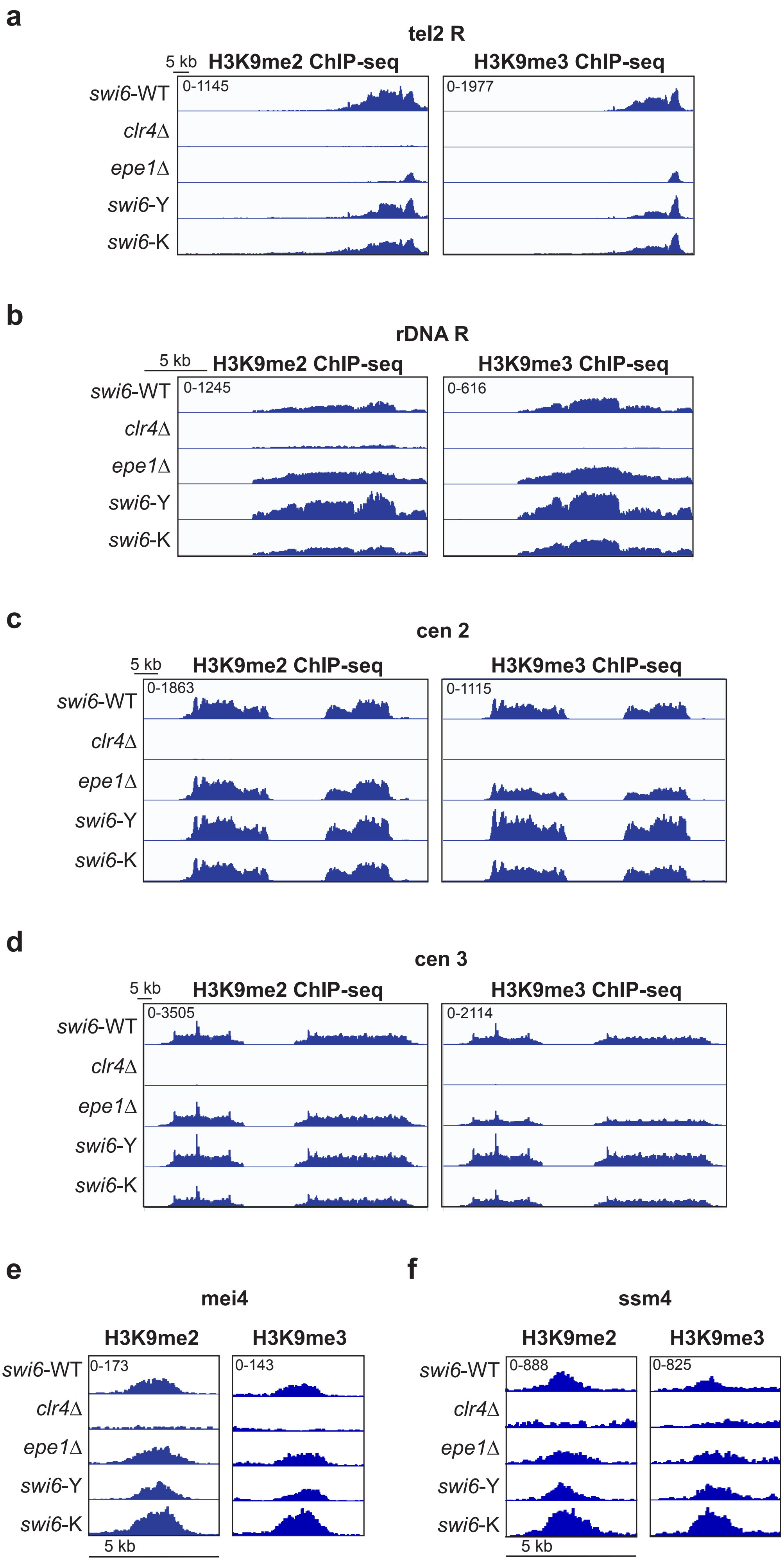
a
tel2 R
5 kb
H3K9me2 ChIP-seq
H3K9me3 ChIP-seq
0-1145
0-1977
swi6-WT
clr4Δ
epe1Δ
swi6-Y
swi6-K
b
rDNA R
5 kb
H3K9me2 ChIP-seq
H3K9me3 ChIP-seq
0-1245
0-616
swi6-WT
clr4Δ
epe1Δ
swi6-Y
swi6-K
c
cen 2
5 kb
H3K9me2 ChIP-seq
H3K9me3 ChIP-seq
0-1863
0-1115
swi6-WT
clr4Δ
epe1Δ
swi6-Y
swi6-K
d
cen 3
5 kb
H3K9me2 ChIP-seq
H3K9me3 ChIP-seq
0-3505
0-2114
swi6-WT
clr4Δ
epe1Δ
swi6-Y
swi6-K
e
mei4
H3K9me2
H3K9me3
0-173
0-143
swi6-WT
clr4Δ
epe1Δ
swi6-Y
swi6-K
5 kb
f
ssm4
H3K9me2
H3K9me3
0-888
0-825
swi6-WT
clr4Δ
epe1Δ
swi6-Y
swi6-K
5 kb

**Supplemental Figure 2. Enrichment of H3K9me2 and H3K9me3 at endogenous heterochromatin sites.** (**a**) right telomere on chromosome II (tel2R). Each panel represents a region spanning 66kb (**b**) rDNA locus on right arm of chromosome III (rDNA-R). Each panel represents a region spanning 19kb. (**c**) centromere on chromosome II and the (**d**) centromere on chromosome III. Each panel represents a region spanning 77 kb region; and the heterochromatin islands at (**e**) *mei4* and (**f**) *ssm4* genes. Each panel represents a region spanning 5 kb. All samples are normalized as reads per kilobase million (RPKM).

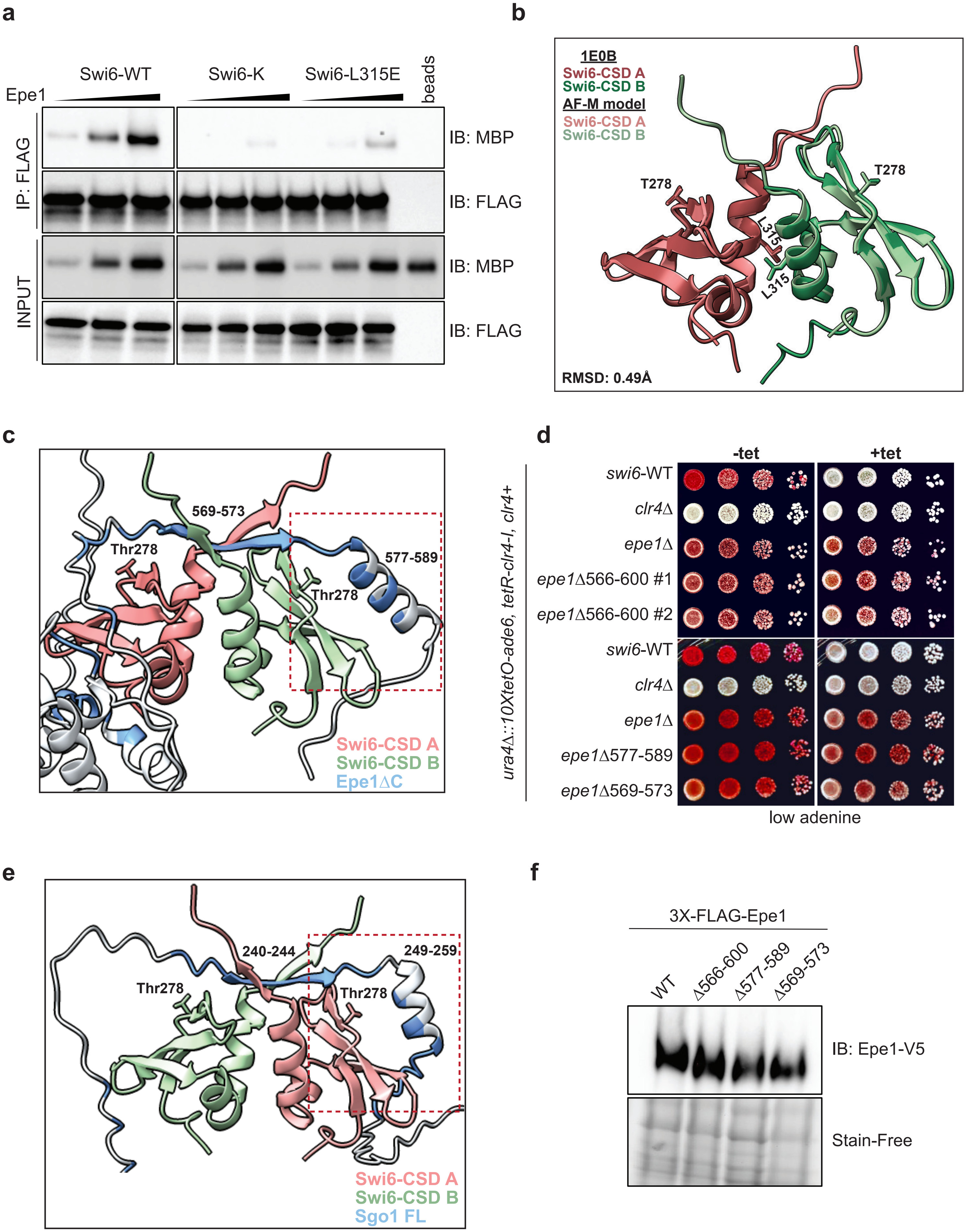

a
Swi6-WT
Swi6-K
Swi6-L315E
beads
Epe1
IP: FLAG
INPUT
IB: MBP
IB: FLAG
IB: MBP
IB: FLAG
b
1E0B
Swi6-CSD A
Swi6-CSD B
AF-M model
Swi6-CSD A
Swi6-CSD B
T278
T278
L315
L315
RMSD: 0.49Å
c
569-573
Thr278
577-589
Thr278
Swi6-CSD A
Swi6-CSD B
Epe1ΔC
d
-tet
+tet
ura4Δ::10XtetO-ade6, tetR-clr4-I, clr4+
swi6-WT
clr4Δ
epe1Δ
epe1Δ566-600 #1
epe1Δ566-600 #2
swi6-WT
clr4Δ
epe1Δ
epe1Δ577-589
epe1Δ569-573
low adenine
e
240-244
249-259
Thr278
Thr278
Swi6-CSD A
Swi6-CSD B
Sgo1 FL
f
3X-FLAG-Epe1
WT
Δ566-600
Δ577-589
Δ569-573
IB: Epe1-V5
Stain-Free

**Supplemental Figure 3. Swi6-Y and Swi6-K perturb Epe1-mediated transcriptional effects through a loss of interaction with Epe1.** (**a**) Western blots of *in vitro* binding assays performed with recombinant 3XFLAG-Swi6-WT, Swi6-K or Swi6 L315E and MBP-Epe1 proteins to test Epe1-Swi6 interaction. Epe1 is detected using an MBP antibody and Swi6 is detected using a FLAG antibody. Background (beads) represents MBP-Epe1 binding non-specifically to M2 FLAG conjugated agarose beads. (**b-c, e**) Highest ranking Alphafold2 Multimer models of (**b)** Swi6-CSD aligned with existing Swi6-CSD crystal structure (PDB 1E0B, 1.9Å). (**c**) Swi6-CSD interaction with Epe1ΔC, encompassing 1-600 amino acid residues. (**e**) Swi6-CSD interaction with meiotic cohesin protection protein, shugoshin, Sgo1. Both structural predictions show a possible interaction between the T278-containing beta sheet interface and a helix downstream of the PxVxL-like motif binding site within the interacting protein. The amino acids predicted by Chimera to form an interface with the Swi6-CSD dimer are shown in blue. Model Confidence parameters and other models reported for Swi6-CSD, Swi6-CSD-Epe1ΔC and Swi6-CSD-Sgo1 are shown in Figures S6-8. (**d**) Silencing assay of *ura4Δ::10XTetO-ade6+* reporter in indicated genotypes in the absence (-tet) and presence (+tet) of tetracycline. Cells are plated at 10-fold serial dilutions. Internal deletions of residues 566-600, 577-589, and 569-573 in Epe1 based on AlphaFold2 Multimer predictions of the interaction between Swi6-CSD-Epe1ΔC. (**f**) Western Blot measuring 3X-FLAG-Epe1 protein levels of full-length (WT) and indicated Epe1 variants. 3X-FLAG-Epe1 is detected using an M2-HRP conjugated antibody and total protein is detected using stain-free imaging.

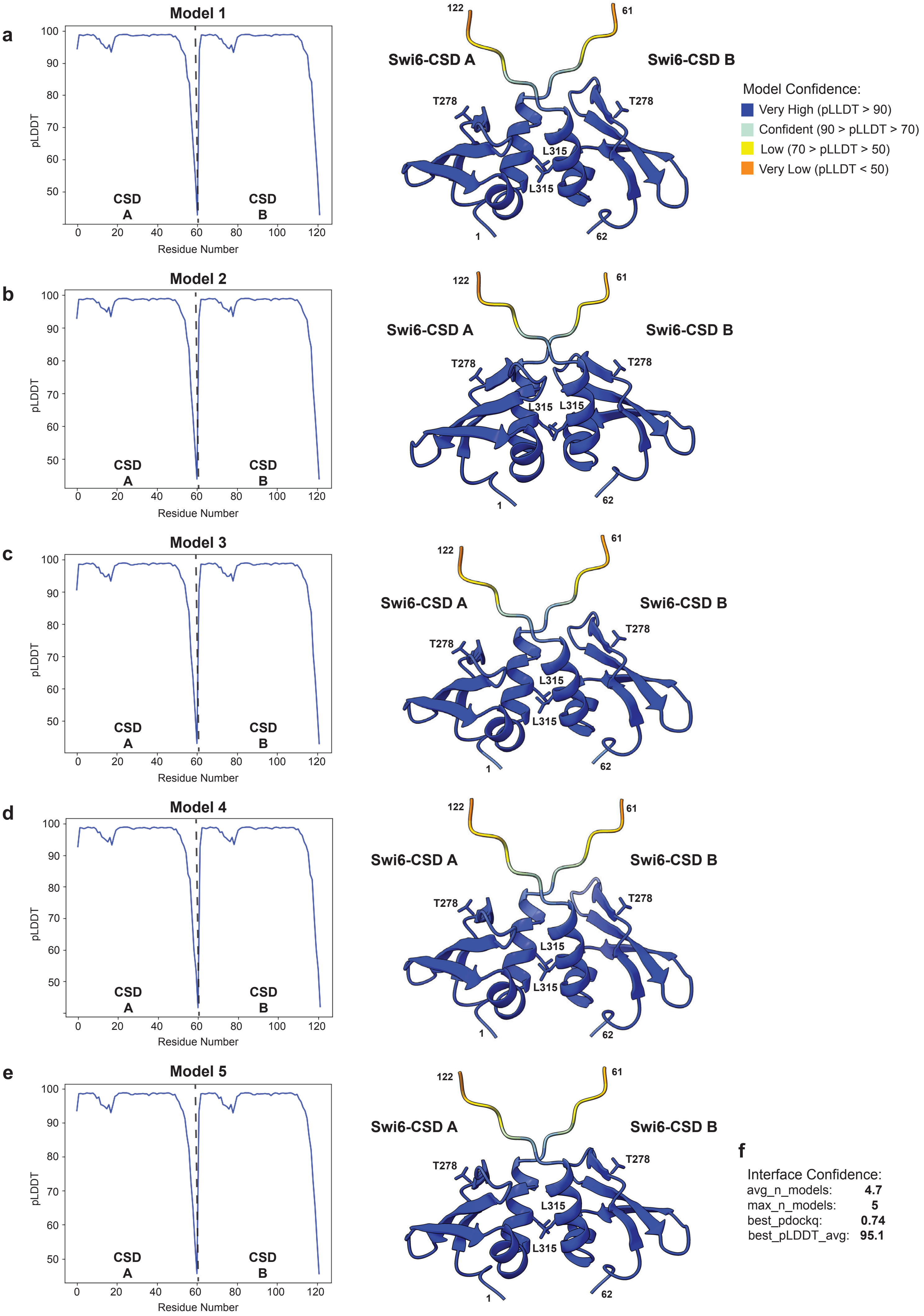

a
Model 1
b
Model 2
c
Model 3
d
Model 4
e
Model 5
pLDDT
Residue Number
CSD A
CSD B
Swi6-CSD A
Swi6-CSD B
T278
L315
122
61
1
62
Model Confidence:
Very High (pLLDT > 90)
Confident (90 > pLLDT > 70)
Low (70 > pLLDT > 50)
Very Low (pLLDT < 50)
f
Interface Confidence:
avg_n_models: 4.7
max_n_models: 5
best_pdockq: 0.74
best_pLDDT_avg: 95.1

**Supplemental Figure 4. Model confidence pLLDT plots and structures for Swi6-CSD dimer predictions generated by AlphaFold2 Multimer.** (**a-e**) Models are ordered in ranking order with (**a**) the highest-ranking structure first and (**e**) the lowest ranking structure last. Structures generated using Chimera and is colored by residue based on the corresponding pLLDT confidence value. The numbering on the structure corresponds to the amino acid position numbering in the pLDDT plot. (**f**) The listed parameters were determined using the code described previously[109]. The avg_n_models parameter quantifies the number of models that predicted the same interface. The max_n_models describes the number of models that satisfy at least some of the contacts at the predicted protein-protein interaction interface. The best_pdockq metric predicts accuracy from 0 (worst) to 1 (best) that considers the average pLDDT of interacting residues and the number of interacting residues, a value of > 0.25 is typical for high confidence predictions. The best_plddt_avg is the calculated average pLDDT of the residues withing the protein-protein interaction interface for the highest-ranking model ranging from 0 (worst) to 100 (best), Values > 70 are typical for confident structure predictions.

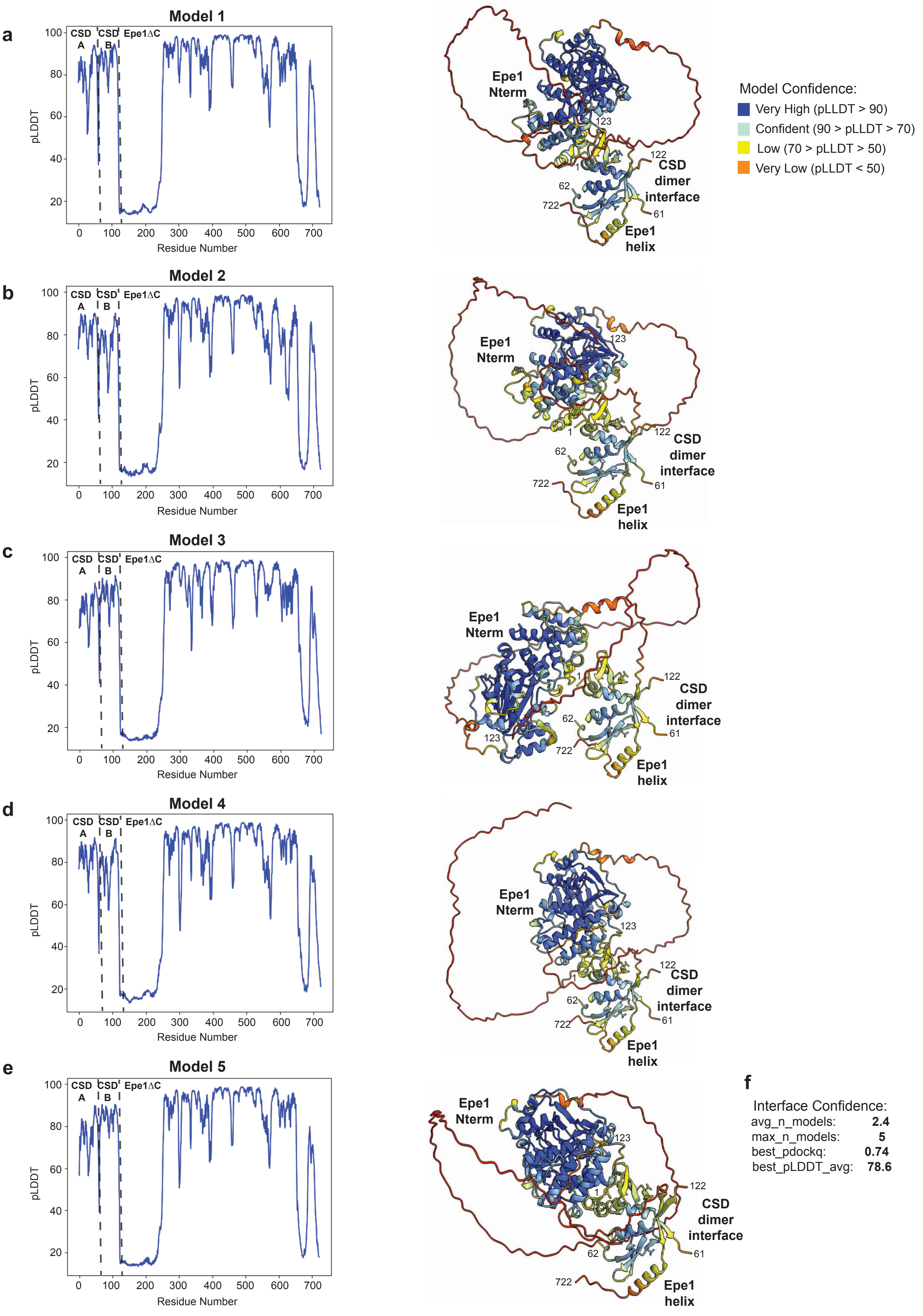
a
Model 1
b
Model 2
c
Model 3
d
Model 4
e
Model 5
CSD A
CSD B
Epe1ΔC
pLDDT
Residue Number
Epe1 Nterm
CSD dimer interface
Epe1 helix
123
122
1
62
722
61
Model Confidence:
Very High (pLLDT > 90)
Confident (90 > pLLDT > 70)
Low (70 > pLLDT > 50)
Very Low (pLLDT < 50)
f
Interface Confidence:
avg_n_models: 2.4
max_n_models: 5
best_pdockq: 0.74
best_pLLDT_avg: 78.6

**Supplemental Figure 5. Model confidence pLLDT plots and structural predictions for Swi6-CSD dimer and Epe1ΔC generated by AlphaFold2 Multimer.** (**a-e**) Models are ordered in ranking order with (**a**) the highest-ranking structure first and (**e**) the lowest ranking structure last. Structures generated using Chimera and is colored by residue based on the corresponding pLLDT confidence value. The numbering on the structure corresponds to the amino acid position numbering in the plot. (**f**) The listed parameters were determined using the code described previously[109]. The avg_n_models parameter quantifies the number of models that predicted the same interface. The max_n_models describes the number of models that satisfy at least some of the contacts at the predicted protein-protein interaction interface. The best_pdockq metric predicts accuracy from 0 (worst) to 1 (best) that considers the average pLDDT of interacting residues and the number of interacting residues, a value of > 0.25 is typical for high confidence predictions. The best_plddt_avg is the calculated average pLDDT of the residues withing the protein-protein interaction interface for the highest-ranking model ranging from 0 (worst) to 100 (best), Values > 70 are typical for confident structure predictions.

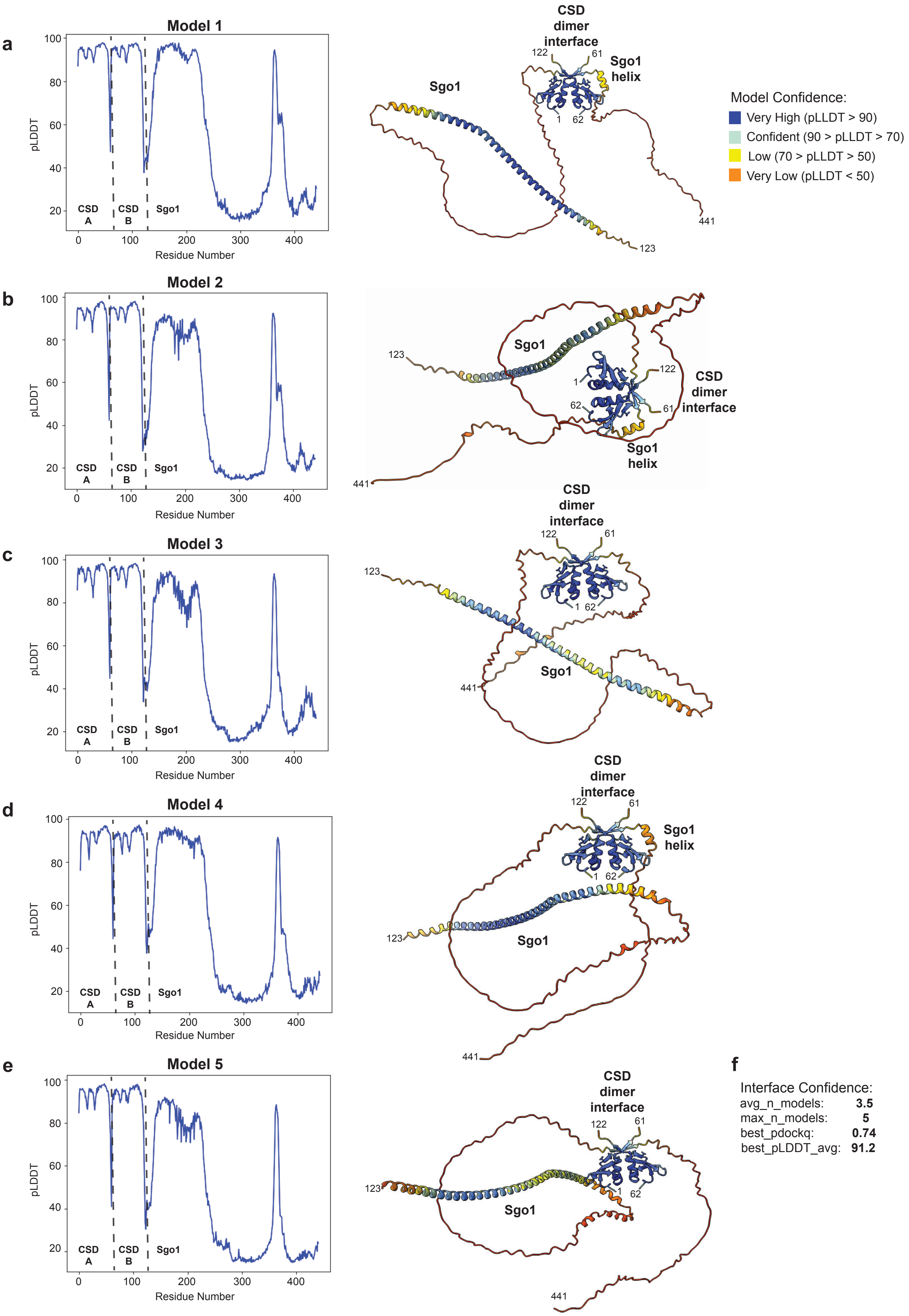
a
Model 1
pLDDT
Residue Number
CSD A
CSD B
Sgo1
CSD dimer interface
Sgo1 helix
Sgo1
122
61
1
62
441
123
Model Confidence:
Very High (pLDDT > 90)
Confident (90 > pLDDT > 70)
Low (70 > pLDDT > 50)
Very Low (pLDDT < 50)
b
Model 2
Sgo1
CSD dimer interface
Sgo1 helix
c
Model 3
d
Model 4
e
Model 5
f
Interface Confidence:
avg_n_models: 3.5
max_n_models: 5
best_pdockq: 0.74
best_pLDDT_avg: 91.2

**Supplemental Figure 6. Model confidence pLLDT plots and structural predictions for Swi6-CSD dimer and Sgo1 generated by AlphaFold2 Multimer.** (**a-e**) Models are ordered in ranking order with (**a**) the highest-ranking structure first and (**e**) the lowest ranking structure last. Structures generated using Chimera and is colored by residue based on the corresponding pLLDT confidence value. The numbering on the structure corresponds to the amino acid position numbering in the plot. (**f**) The listed parameters were determined using the code described previously[109]. The avg_n_models parameter quantifies the number of models that predicted the same interface. The max_n_models describes the number of models that satisfy at least some of the contacts at the predicted protein-protein interaction interface. The best_pdockq metric predicts accuracy from 0 (worst) to 1 (best) that considers the average pLDDT of interacting residues and the number of interacting residues, a value of > 0.25 is typical for high confidence predictions. The best_plddt_avg is the calculated average pLDDT of the residues withing the protein-protein interaction interface for the highest-ranking model ranging from 0 (worst) to 100 (best), Values > 70 are typical for confident structure predictions.

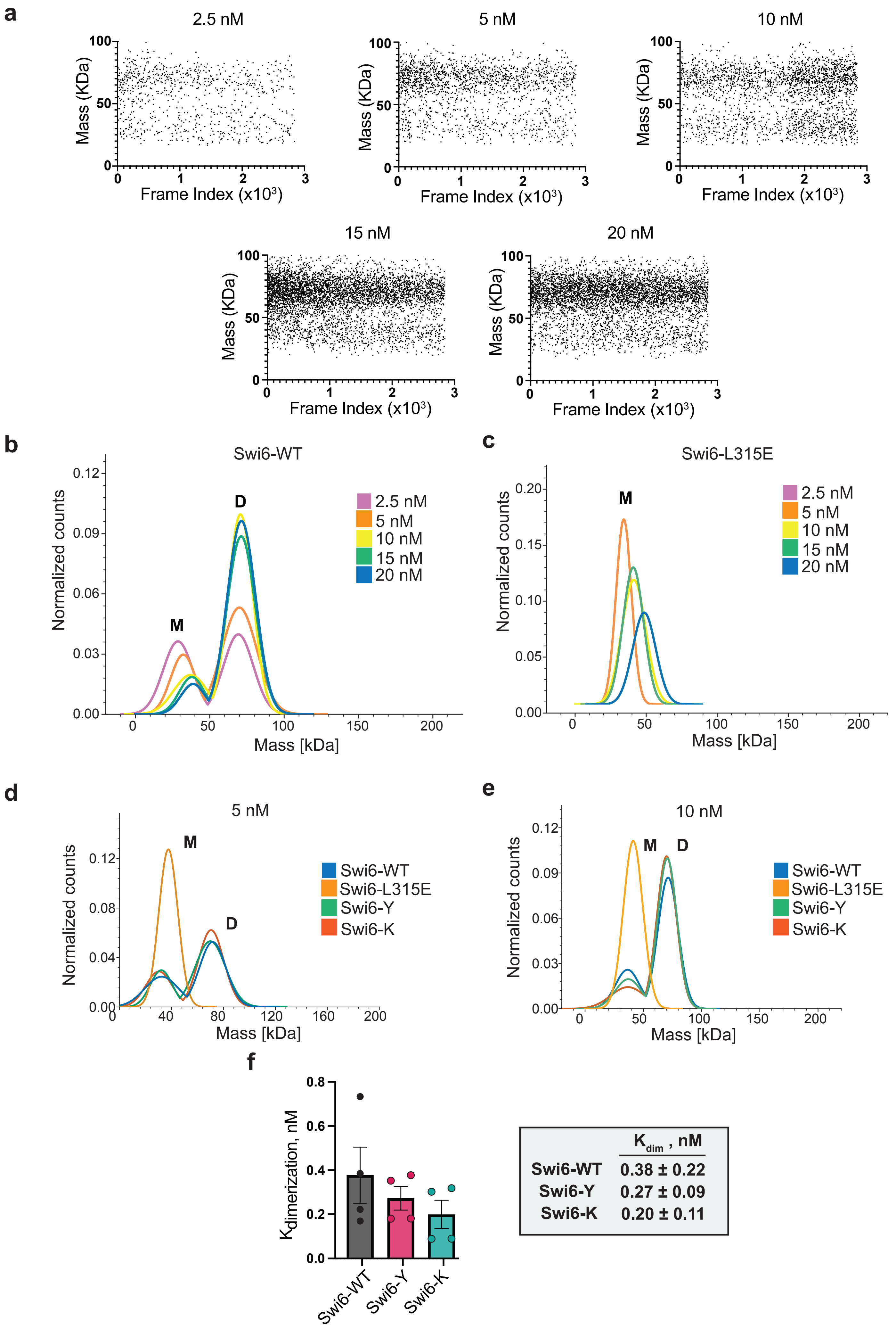
a
2.5 nM
5 nM
10 nM
15 nM
20 nM
Mass (KDa)
Frame Index (x10^3)
b
Swi6-WT
Normalized counts
Mass [kDa]
M
D
2.5 nM
5 nM
10 nM
15 nM
20 nM
c
Swi6-L315E
d
5 nM
Swi6-WT
Swi6-L315E
Swi6-Y
Swi6-K
e
10 nM
f
Kdimerization, nM
Swi6-WT
Swi6-Y
Swi6-K
Kdim , nM
Swi6-WT 0.38 ± 0.22
Swi6-Y 0.27 ± 0.09
Swi6-K 0.20 ± 0.11

**Supplemental Figure 7. Relative abundance of Swi6 monomer and dimer molecular species detected by mass photometry with increasing Swi6 concentration**. (**a**) Density plot of detected molecules of indicated masses as a function of time (frame index) across different Swi6 concentrations. (**b-c**) Representative gaussian distributions of (**b**) Swi6-WT and (**c**) Swi6 L315E. Monomers (M) and dimers (D) are marked based on expected molecular weights of Swi6. (**d-e**) Representative gaussian distributions for indicated Swi6 proteins at (**d**) 5 nM and (**e**) 10 nM concentrations. (**f**) Left- Bar graph of the mean Swi6 dimerization affinity ($K_{dim}$) calculated by molecular counting across replicates, error bars indicate SD (N= 4). Right- Table summarizing Swi6 $K_{dim}$ values. Error bars represent SD (N=4).

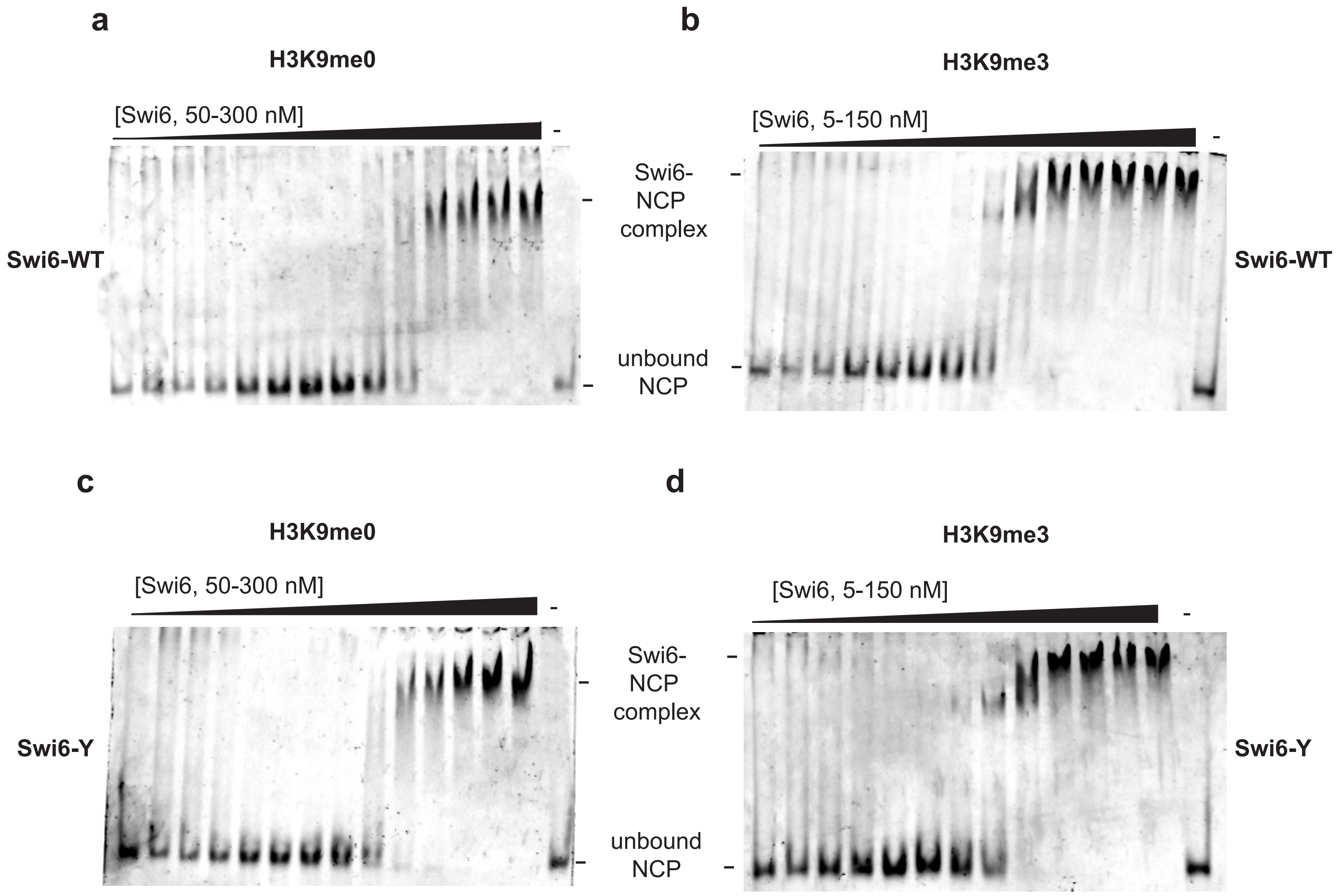
a
H3K9me0
[Swi6, 50-300 nM]
-
Swi6-WT
Swi6-
NCP
complex
unbound
NCP
b
H3K9me3
[Swi6, 5-150 nM]
-
Swi6-WT
c
H3K9me0
[Swi6, 50-300 nM]
-
Swi6-Y
Swi6-
NCP
complex
unbound
NCP
d
H3K9me3
[Swi6, 5-150 nM]
-
Swi6-Y

**Supplemental Figure 8. Analysis of Swi6 nucleosome binding *in vitro* using EMSAs.** (**a-d**) Representative images of (**a,c**) H3K9me0 mononucleosome and (**b,d**) H3K9me3 mononucleosome gel shifts with increasing (**a,b**) Swi6-WT and (**c,d**) Swi6-Y concentrations. The lower band represents unbound, intact nucleosome core particles (NCPs) and the disappearance of that band indicates binding of the NCP to Swi6, accompanied by the appearance of higher molecular weight species.

**a**

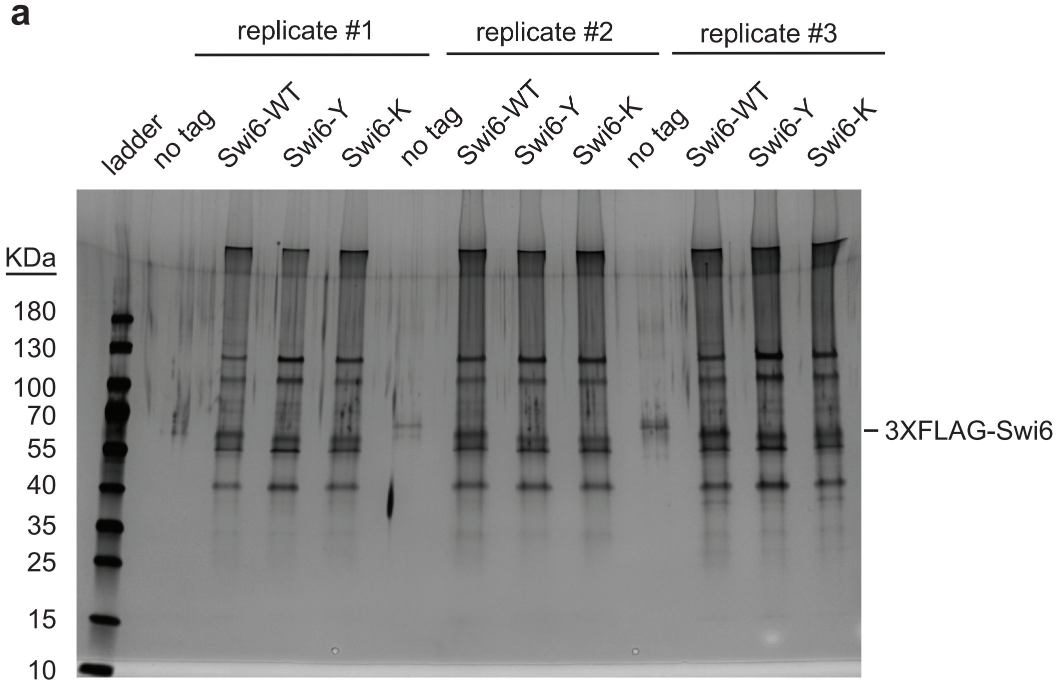


**b**

**Known heterchromatin maintenance regulators without detectable enrichment in Swi6-Y/K**

**CLRC:** Clr4, Raf1, Raf2, Rik1, Cul4, Rbx1
**SHREC:** Chp2, Mit1, Clr3, Clr2
**FACT:** Pob3, Spt16
**NPC:** Alm1, Amo1, Npp106, Lem2
**Ino80:** Iec5

not detected by TMT-MS (Cul4, Rbx1, Alm1, Amo1)
no significant enrichment over Swi6-wt

**c**

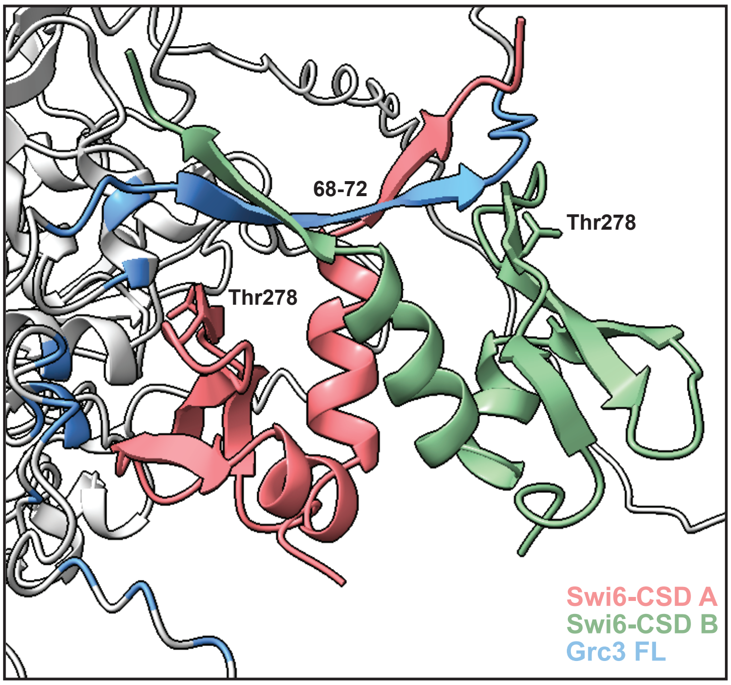

**Supplemental Figure 9. Quantitative proteomics reveal specific enrichment of the rixosome complex with Swi6-Y.** (**a**) Silver stain of 3XFLAG-Swi6 purification samples used for TMT-MS analysis (N=3). (**b**) Proteins that have established roles in heterochromatin maintenance that did not have significant enrichment compared to Swi6-WT (blue) or were not detected (red) during TMT-MS. (**c**) Highest ranking Alphafold2 Multimer model predicting Swi6-CSD dimer interaction with Grc3. The model reveals a PxVxL motif in Grc3 interacting with the Swi6-CSD dimer interface but unlike the models presented earlier, the downstream region is disordered and does not show a predicted interaction with the Thr 278-containing beta sheet interface. The amino acids predicted by Chimera to form an interface with the Swi6-CSD dimer are shown in blue.

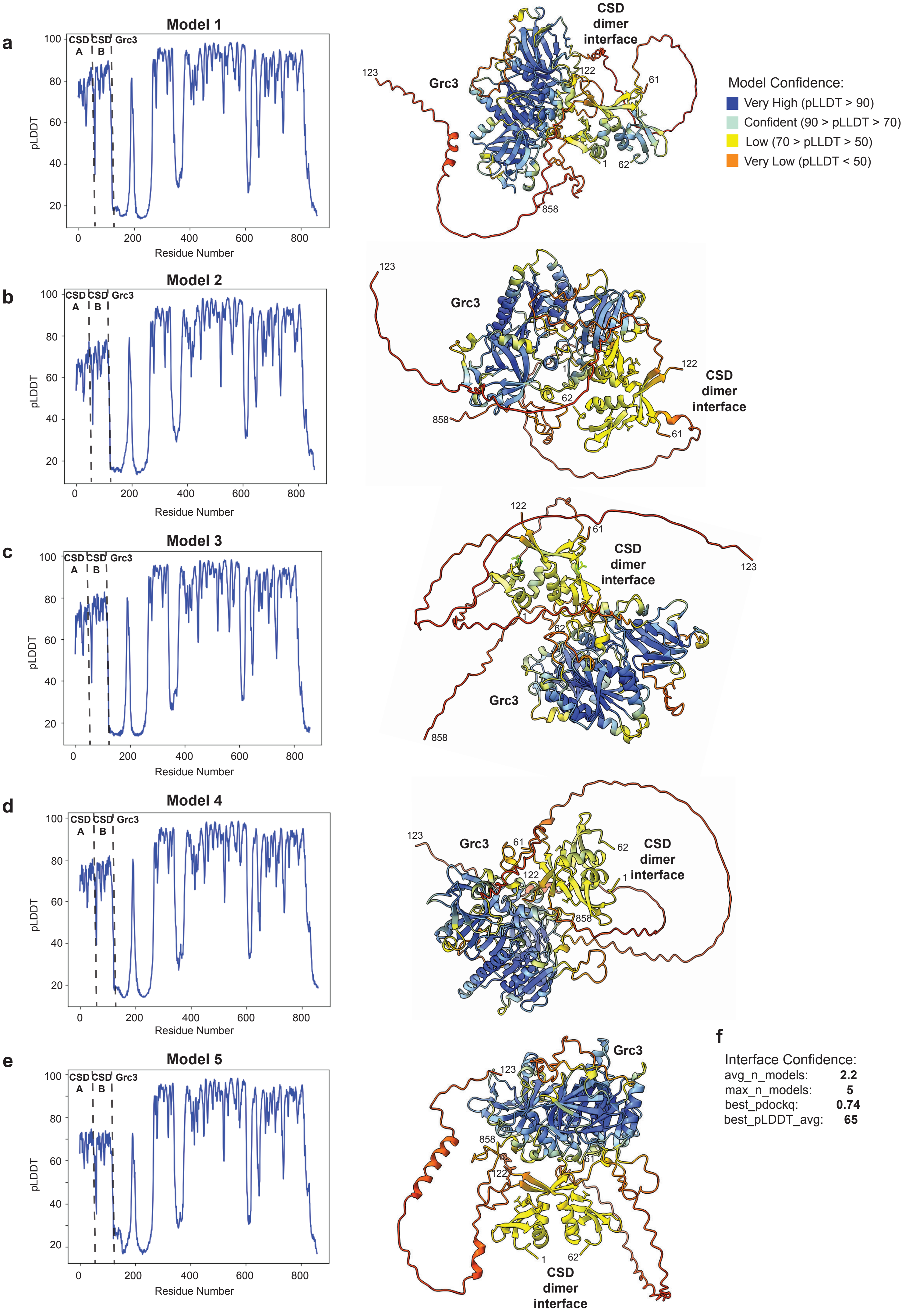

a
Model 1
b
Model 2
c
Model 3
d
Model 4
e
Model 5
CSD A
CSD B
Grc3
pLDDT
Residue Number
CSD dimer interface
Model Confidence:
Very High (pLLDT > 90)
Confident (90 > pLLDT > 70)
Low (70 > pLLDT > 50)
Very Low (pLLDT < 50)
f
Interface Confidence:
avg_n_models: 2.2
max_n_models: 5
best_pdockq: 0.74
best_pLDDT_avg: 65

**Supplemental Figure 10. Model confidence pLLDT plots and structural predictions for Swi6-CSD dimer and Grc3 generated by AlphaFold2 Multimer.** (**a-e**) Models are ordered in ranking order with (**a**) the highest-ranking structure first and (**e**) the lowest ranking structure last. Structures generated using Chimera and is colored by residue based on the corresponding pLLDT confidence value. The numbering on the structure corresponds to the amino acid position numbering in the plot. (**f**) The listed parameters were determined using the code described previously[109]. The avg_n_models parameter quantifies the number of models that predicted the same interface. The max_n_models describes the number of models that satisfy at least some of the contacts at the predicted protein-protein interaction interface. The best_pdockq metric predicts accuracy from 0 (worst) to 1 (best) that considers the average pLDDT of interacting residues and the number of interacting residues, a value of > 0.25 is typical for high confidence predictions. The best_plddt_avg is the calculated average pLDDT of the residues withing the protein-protein interaction interface for the highest-ranking model ranging from 0 (worst) to 100 (best), Values > 70 are typical for confident structure predictions.

**a**

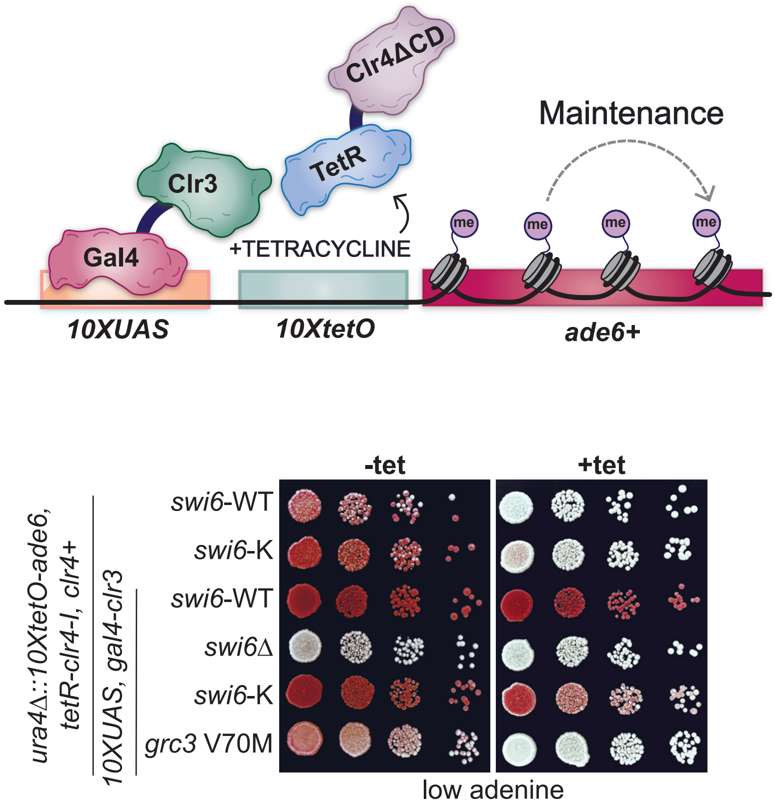
Clr4ΔCD
TetR
Clr3
Gal4
+TETRACYCLINE
Maintenance
me
10XUAS
10XtetO
ade6+
-tet
+tet
ura4Δ::10XtetO-ade6, tetR-clr4-I, clr4+
10XUAS, gal4-clr3
swi6-WT
swi6-K
swi6-WT
swi6Δ
swi6-K
grc3 V70M
low adenine

**b**

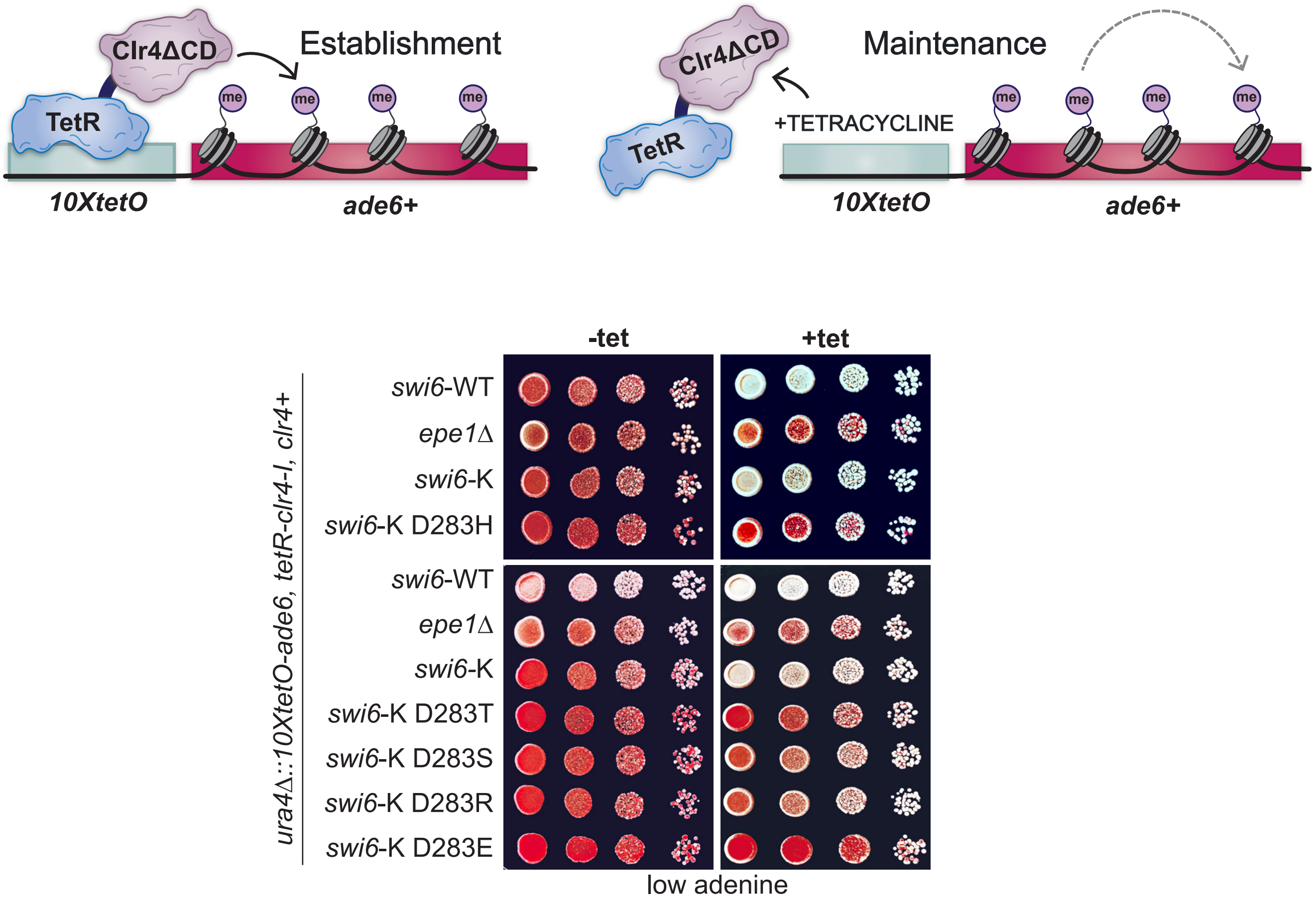
Clr4ΔCD
TetR
Establishment
me
10XtetO
ade6+
Clr4ΔCD
TetR
+TETRACYCLINE
Maintenance
10XtetO
ade6+
-tet
+tet
ura4Δ::10XtetO-ade6, tetR-clr4-I, clr4+
swi6-WT
epe1Δ
swi6-K
swi6-K D283H
swi6-WT
epe1Δ
swi6-K
swi6-K D283T
swi6-K D283S
swi6-K D283R
swi6-K D283E
low adenine

**Supplementary Figure 11. The maintenance defect observed in Swi6-K can be genetically rescued.** (**a**) Top- schematic of the modified *10xUAS-10XtetO-ade6+* reporter system. The system contains ten copies of the *UAS* DNA binding sites placed upstream of *10XtetO*. A Gal4-Clr3 fusion protein binds to these sites and remains bound in the presence of tetracycline. Bottom-Silencing assay of *ura4Δ::10xUAS-10XtetO-ade6+* reporter in indicated genotypes in the absence (-tet) and presence (+tet) of tetracycline. Red cells indicate *ade6+* silencing. Cells are plated at 10-fold serial dilutions. (**b**) Top- schematic of the *10XtetO-ade6+* reporter system. Bottom-Silencing assay of *ura4Δ::10XtetO-ade6+* reporter in indicated genotypes in the absence (-tet) and presence (+tet) of tetracycline. Red cells indicate ade6+ silencing. Cells are plated at 10-fold serial dilutions.

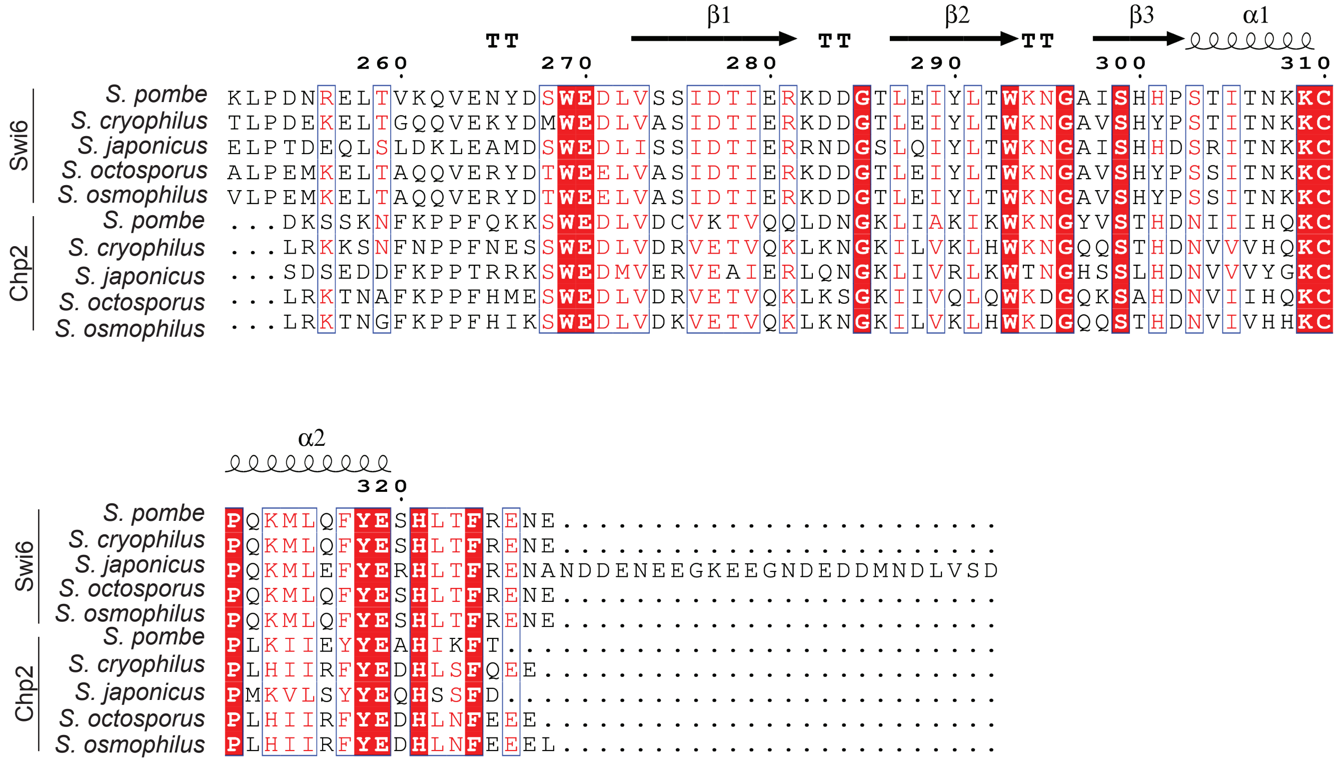

β1
β2
β3
α1
α2
Swi6
Chp2
S. pombe
S. cryophilus
S. japonicus
S. octosporus
S. osmophilus

**Supplemental Figure 12. Multiple sequence alignment (MSA) of Swi6 and Chp2 within the Schizosaccharomyces lineage.** Multiple Sequence Alignment generated using Clustal Omega and plotted using ESPript3.0 of Swi6-CSD (amino acid residues 269-324) or Chp2-CSD (amino acid residues 324-379) from indicated species. Known structural features are plotted above each alignment; residue similarity is indicated as follows: strict identity (red box white character), similarity in a group (red text), similarity across groups (blue frame).

**Supplementary Table 1*.* *S.pombe* strains used in this study.**

| Strain no. | Strain genotype | Source | Related to |
|---|---|---|---|
| KR18 | *h- leu1-32 ade6-M210 ura4Δ::10XtetO-ade6 clr4+ trp1:nat-clr4p-tetR-clr4ΔCD* | Moazed Lab | Figures 1B-G, 2A-B,2E, 3A-B, 5F Supplementary Figures 1B-D, 2, 3D, 11A-B |
| KR24 | *h- leu1-32 ade6-M210 ura4Δ::10XtetO-ade6, clr4Δ::kanMX6* | Moazed Lab | Figures 1B-C,1E-G, 2A-B,2E, 3A-B Supplementary Figures 1D, 2, 3D |
| KR33 | *h- leu1-32 ade6-M210 ura4Δ::10XtetO-ade6 clr4+ trp1:nat-clr4p-tetR-clr4ΔCD epe1Δ::kanMX6* | Moazed Lab | Figures 1B-G, 2A-B,E, 3A-B, 5E Supplementary Figures 1D, 2, 3D, 11A-B |
| KR283 | *h- leu1-32 ade6? ura4Δ::10XtetO-ade6 clr4+, trp1:nat-clr4p-tetR-clr4ΔCD , epe1Δ::3XFLAG-epe1-hph* | Moazed Lab | Supplementary Figure 3F |
| KR343 | *h+ otr1R(SphI)::ade6+ ura4-D18 leu1-32 ade6-M210* | Moazed Lab | Figures 2C-D, 3D, 5A-B, Supplementary Figure 9A |
| KR453 | *h+ otr1R(SphI)::ade6+ ura4-D18 leu1-32 ade6-M210 swi6D::ura4 3XFLAG-Swi6* | Moazed Lab | Figure 5A-C Supplementary Figure 9A |
| KR555 | *h+ otr1R(SphI)::ade6+ ura4-D18 leu1-32 ade6-M210, clr4D::ura4-kanMX6* | Moazed Lab | Figure 2C-D |
| KR778 | *h90 leu1-32 ade6M216 ura4D-18 swi6: PAmCherry-Swi6* | This study | Figure 4E-G |
| KR1101 | *h- leu1-32 ade6-M210 tandem ura4Δ::10XUAS-10XtetO-ade6 natMX6-tetR-clr4-I epe1+ leu1:nmt1-Gal4-Clr3 swi6Δ::* | This study | Supplementary Figure 11A |
| KR1124 | *h+ leu1-32 ura4Δ::10XUAS-tetO-ade6+ natMX6-tetR-clr4-I epe1+ leu1:nmt1-gal4-clr3* | This study | Supplementary Figure 11A |
| KR1982 | *h- leu1-32 ade6-M210 ura4Δ::10XtetO-ade6 clr4+ trp1:nat-clr4p-tetR-clr4ΔCD swi6T278S* | This study | Figure 1B Supplementary Figure 1B |
| KR1984 | *h- leu1-32 ade6-M210 ura4Δ::10XtetO-ade6 clr4+ trp1:nat-clr4p-tetR-clr4ΔCD swi6T278Y* | This study | Figure 1B,1D-G, 2A-B,E, 3A-B, 5E Supplementary Figure 1B-C |
| KR1986 | *h- leu1-32 ade6-M210 ura4Δ::10XtetO-ade6 clr4+ trp1:nat-clr4p-tetR-clr4ΔCD swi6T278F* | This study | Figure 1B Supplementary Figure 1B |
| KR1988 | *h- leu1-32 ade6-M210 ura4Δ::10XtetO-ade6 clr4+ trp1:nat-clr4p-tetR-clr4ΔCD swi6T278C* | This study | Figure 1B Supplementary Figure 1B |
| KR1990 | *h- leu1-32 ade6-M210 ura4Δ::10XtetO-ade6 clr4+ trp1:nat-clr4p-tetR-clr4ΔCD swi6T278K* | This study | Figure 1C-F, 2A-B, 2E, 3A-B Supplementary Figure 1B-C, 11A-B |

| | | | |
|---|---|---|---|
| KR1994 | *h- leu1-32 ade6-M210 ura4Δ::10XtetO-ade6 clr4+ trp1:nat-clr4p-tetR-clr4ΔCD epe1-V5-kanMX6* | This study | Figure 3D |
| KR1997 | *h- leu1-32 ade6-M210 ura4Δ::10XtetO-ade6 clr4+ trp1:nat-clr4p-tetR-clr4ΔCD swi6T278Y epe1Δ::kanMX6* | This study | Figure 1F, 3A-B Supplementary Figure 1D |
| KR1999 | *h- leu1-32 ade6-M210 ura4Δ::10XtetO-ade6 clr4+ trp1:nat-clr4p-tetR-clr4ΔCD ckb1D::kanMX6* | This study | Figure 1G |
| KR2000 | *h- leu1-32 ade6-M210 ura4Δ::10XtetO-ade6 clr4+ trp1:nat-clr4p-tetR-clr4ΔCD swi6T278Y ckb1Δ::kanMX6* | This study | Figure 1G |
| KR2003 | *h+ otr1R(SphI)::ade6+ ura4-D18 leu1-32 ade6-M210 swi6Δ::ura4 3XFLAG-swi6T278Y* | This study | Figure 5A, C Supplementary Figure 9A |
| KR2009 | *h+ otr1R(SphI)::ade6+ ura4-D18 leu1-32 ade6-M210 swi6T278Y* | This study | Figure 2C-D |
| KR2041 | *h+ otr1R(SphI)::ade6+ ura4-D18 leu1-32 ade6-M210 swi6T278K* | This study | Figure 2C-D |
| KR2101 | *h- leu1-32 ade6-M210 ura4Δ::10XtetO-ade6 clr4+ trp1:nat-clr4p-tetR-clr4ΔCD swi6Δ270-328::ura4-hphMX6* | This study | Supplementary Figure 1A |
| KR2436 | *h- leu1-32 ade6-M210 ura4Δ::10XtetO-ade6 clr4+ trp1:nat-clr4p-tetR-clr4ΔCD crb3-TAP-kanMX6* | This study | Figure 5F |
| KR2441 | *h- leu1-32 ade6-M210 ura4Δ::10XtetO-ade6 clr4+ trp1:nat-clr4p-tetR-clr4DΔCD epe1-V5-hphMX6 swi6T278Y* | This study | Figure 3D |
| KR2443 | *h- leu1-32 ade6-M210 ura4Δ::10XtetO-ade6 clr4+ trp1:nat-clr4p-tetR-clr4ΔCD epe1-V5-hphMX6 swi6T278K* | This study | Figure 3D |
| KR2445 | *h90 leu1-32 ade6M216 ura4D-18 swi6: PAmCherry-swi6T278K* | This study | Figure 4F-G |
| KR2465 | *h- leu1-32 ade6-M210 ura4Δ::10XtetO-ade6 trp1:nat-clr4p-tetR-clr4ΔCD swi6T278Y grc3V70M* | This study | Figure 5E |
| KR2468 | *h- leu1-32 ade6-M210 ura4Δ::10XtetO-ade6 trp1:nat-clr4p-tetR-clr4ΔCD epe1Δ-kanMX6 grc3V70M* | This study | Figure 5E |
| KR2471 | *h90 leu1-32 ade6M216 ura4D-18 swi6: PAmCherry-swi6T278Y* | This study | Figure 4E, G |

| | | | |
|---|---|---|---|
| KR2474 | *h- leu1-32 ade6-M210 ura4Δ::10XtetO-ade6 clr4+ trp1:nat-clr4p-tetR-clr4ΔCD swi6T278Y crb3-TAP-kanMX6* | This study | Figure 5F |
| KR2650 | *h- leu1-32 ade6-M210 ura4Δ::10XtetO-ade6 clr4+ trp1:nat-clr4p-tetR-clr4ΔCD swi6T278K epe1Δ::kanMX6* | This study | Figure 1F, 3A-B Supplementary Figure 1D |
| KR2686 | *h- leu1-32 ade6-M210 ura4Δ::10XUAS-tetO-ade6 natMX6-tetR-clr4-I epe1+ leu1:nmt1-gal4-clr3 swi6T278K* | This study | Supplementary Figure 11A |
| KR2700 | *h- leu1-32 ade6-M210 ura4Δ::10XtetO-ade6 clr4+ trp1:nat-clr4p-tetR-clr4ΔCD swi6T278K crb3-TAP-kanMX6* | This study | Figure 5F |
| KR2703 | *h- leu1-32 ade6-M210 ura4Δ::10XtetO-ade6 clr4+ trp1:nat-clr4p-tetR-clr4ΔCD grc3V70M crb3-TAP-kanMX6* | This study | Figure 5F |
| KR2736 | *h- leu1-32 ade6-M210 ura4Δ::10XtetO-ade6 clr4+ trp1:nat-clr4p-tetR-clr4ΔCD swi6T278R* | This study | Figure 1C Supplementary Figure 1B |
| KR2798 | *h- leu1-32 ade6-M210 ura4Δ::10XtetO-ade6 clr4+ trp1:nat-clr4p-tetR-clr4ΔCD swi6T278A* | This study | Figure 1B Supplementary Figure 1B |
| KR2799 | *h- leu1-32 ade6-M210 ura4Δ::10XtetO-ade6 clr4+ trp1:nat-clr4p-tetR-clr4ΔCD swi6T278D* | This study | Figure 1C Supplementary Figure 1B |
| KR2801 | *h- leu1-32 ade6-M210 ura4Δ::10XtetO-ade6 clr4+ trp1:nat-clr4p-tetR-clr4ΔCD swi6T278E* | This study | Figure 1C Supplementary Figure 1B |
| KR2825 | *h+ otr1R(SphI)::ade6+ ura4-D18 leu1-32 ade6-M210 swi6Δ::ura4 3XFLAG-swi6T278K* | This study | Figure 5B-C Supplementary Figure 9A |
| KR2827 | *h- leu1-32 ade6-M210 ura4Δ::10XtetO-ade6 clr4+ trp1:nat-clr4p-tetR-clr4ΔCD ckb1Δ::kanMX6 epe1Δ::hphMX6* | This study | Figure 1G |
| KR2867 | *h+ leu1-32 ura4Δ::10XUAS-tetO-ade6 natMX6-tetR-clr4-I epe1+ leu1:nmt1-gal4-clr3 grc3V70M* | This study | Supplementary Figure 11A |
| KR3086 | *h- leu1-32 ade6? ura4Δ::10XtetO-ade6 clr4+, trp1:nat-clr4p-tetR-clr4ΔCD , epe1Δ::3XFLAG-epe1Δ566-600-hph* #1 | This study | Supplementary Figure 3D,F |
| KR3087 | *h- leu1-32 ade6? ura4Δ::10XtetO-ade6 clr4+, trp1:nat-clr4p-tetR-clr4ΔCD , epe1Δ::3XFLAG-epe1Δ566-600-hph* #2 | This study | Supplementary Figure 3D |

| | | | |
|---|---|---|---|
| KR3127 | *h- leu1-32 ade6? ura4∆::10XtetO-ade6 clr4+, trp1:nat-clr4p-tetR-clr4∆CD , epe1∆::3XFLAG-epe1∆569-573-hph* | This study | Supplementary Figure 3D,F |
| KR3167 | *h- leu1-32 ade6? ura4∆::10XtetO-ade6 clr4+, trp1:nat-clr4p-tetR-clr4∆CD , epe1∆::3XFLAG-epe1∆577-589-hph* | This study | Supplementary Figure 3D,F |
| KR3284 | *h- leu1-32 ade6-M210 ura4∆::10XtetO-ade6 clr4+ trp1:nat-clr4p-tetR-clr4∆CD swi6T278KD283H* | This study | Supplementary Figure 11B |
| KR3285 | *h- leu1-32 ade6-M210 ura4∆::10XtetO-ade6 clr4+ trp1:nat-clr4p-tetR-clr4∆CD swi6T278KD283T* | This study | Supplementary Figure 11B |
| KR3286 | *h- leu1-32 ade6-M210 ura4∆::10XtetO-ade6 clr4+ trp1:nat-clr4p-tetR-clr4∆CD swi6T278KD283S* | This study | Supplementary Figure 11B |
| KR3287 | *h- leu1-32 ade6-M210 ura4∆::10XtetO-ade6 clr4+ trp1:nat-clr4p-tetR-clr4∆CD swi6T278KD283R* | This study | Supplementary Figure 11B |
| KR3288 | *h- leu1-32 ade6-M210 ura4∆::10XtetO-ade6 clr4+ trp1:nat-clr4p-tetR-clr4∆CD swi6T278KD283E* | This study | Supplementary Figure 11B |

**Supplementary Table 2. Primers used in this study for quantitative PCR**

| Primer | Sequence |
|---|---|
| *SPCC330.06c F* | GCCGTAAATGACGTTTTCGTCACC |
| *SPCC330.06c R* | ACCTTGACAACCTTGCCATTCTCG |
| *tub F* | AACGCTTGGCCATGGAATACACG |
| *tub R* | GAGAGGCGGTGATGGAAGAAACAAC |
| *dg F* | GCGGTCATTTAAAGGCATAG |
| *dg R* | CGACAACACGAATTTTACTTCTAGGTC |
| *dh F* | GTATTTGGATTCCATCGGTACTATGG |
| *dh R* | ACTACATCGACACAGAAAAGAAAACAA |
| *tlh1 F* | AGAAACACCGAAACCAGCCA |
| *tlh1 R* | TGGCGATACAGACGAAGGGA |
